# Non-destructive Chemical Imaging of Bone Tissue for Intraoperative and Diagnostic Applications

**DOI:** 10.1101/2021.05.14.444201

**Authors:** Kseniya S. Shin, Shuaiqian Men, Angel Wong, Colburn Cobb-Bruno, Eleanor Chen, Dan Fu

## Abstract

Bone is difficult to image using traditional histopathological methods, leading to challenges in intraoperative consultations needed in orthopedic oncology. However, intraoperative pathological evaluation is critical in guiding surgical treatment. In this study, we demonstrate that a multimodal imaging approach that combines stimulated Raman scattering (SRS) microscopy, two-photon fluorescence (TPF) microscopy, and second harmonic generation (SHG) microscopy can provide useful diagnostic information regarding intact bone tissue fragments from surgical excision or biopsy specimens. We imaged bone samples from 14 patient cases and performed comprehensive chemical and morphological analyses of both mineral and organic components of bone. Our main findings show that carbonate content combined with morphometric analysis of bone organic matrix can separate several major classes of bone cancer associated diagnostic categories with an average accuracy of >90%. This proof-of-principle study demonstrate that multimodal imaging and machine learning-based analysis of bony tissue can provide crucial diagnostic information for guiding clinical decisions in orthopedic oncology.

## Introduction

Intraoperative consultation of bone specimens plays an important role in orthopedic surgery. One of the main reasons for intraoperative consultation on a bone specimen is to determine whether an intra-osseous lesion is benign or malignant (*1*). In the case of a malignant lesion, separating primary bone cancer from metastasis becomes another challenge (*2*). An accurate intraoperative diagnosis guides subsequent clinical decisions on operative procedures and clinical management.

Current intraoperative pathology consultations rely heavily on gross examination, cytologic preparations, and frozen sections. While these approaches work well for many organ systems, bone samples have proved to be challenging due to their mineralized content. Sectioning-dependent techniques are not feasible for histologic evaluation in an intraoperative setting, as de-mineralization protocols for bone specimens can take up to one week. Additionally, the de-mineralization procedure destroys the DNA/RNA necessary for clinical molecular tests and compromises the histologic quality of tissue sections, making definitive pathologic diagnosis difficult (*3*). Although soft tissue and relevant cellular materials can sometimes supplement and guide medical treatment of musculoskeletal lesions (*4, 5*), they are often insufficient for making clinical decisions. Without the ability to interrogate bone tissue intraoperatively, the patient must wait for post-surgical tissue processing and possibly undergo additional procedures based on pending pathology diagnosis. Thus, there is an unmet need to develop tools capable of providing intraoperative diagnosis of bone specimens to better inform surgeons as they decide on surgical treatment.

Bone is an important specialized connective tissue composed of mineralized extracellular material (primarily type I collagen). The mineral content of bone consists mainly of hydroxyapatite [Ca_10_(PO_4_)_6_(OH)_2_], with variable amounts of carbonate and magnesium (*6*). Conventional light microscopy relies on H&E staining for morphology and various special stains (e.g., Masson’s trichrome and von Kossa) to visualize collagen and mineralization. Special stains require time-consuming processing procedures, which is why such approaches are only used for permanent processing of specimens after surgery. To visualize the internal structures of bone specimens, several ultrastructure-geared techniques including scanning electron microscopy and small-angle x-ray scattering microscopy have been used for characterization. Both techniques are time consuming and destructive and thus are inapplicable to timely clinical diagnosis.

Optical techniques such as second harmonic generation (SHG) microscopy (*7*), third harmonic generation (THG) microscopy (*8*), Fourier-transform infrared spectroscopy (FTIR) (*9*), and Raman spectroscopy (*10*–*14*) allow for noninvasive qualitative and quantitative evaluation of bone specimens. These techniques have been used to study mostly non-neoplastic processes of bone including healing, aging, osteoporosis, and osteomyelitis (*12, 15*–*18*). A few reports have probed chemical changes of mineral components occurring in a very limited collection of cancers (*19, 20*). Nonetheless, our understanding of chemical changes in bone related to primary neoplastic processes such as osteosarcoma and chondrosarcoma remains limited. Moreover, there has been no attempt to explore chemical changes for assessing pathological conditions of bone specifically for intraoperative applications.

Stimulated Raman scattering (SRS) microscopy, a powerful label-free technique capable of providing chemical information at submicron spatial resolution (*21*), has shown tremendous promise for label-free histopathology applications. One emerging variation is Stimulated Raman histology (SRH), which works by exploiting lipid and protein Raman contrasts to generate H&E equivalent images (*21, 22*). Applications of SRH to diagnosis of brain tumors (*22*–*24*), laryngeal cancer (*25*), gastrointestinal cancer (*26*) demonstrate the promise of SRH as an alternative to H&E. However, the lack of contrast between bone matrix and typical cellular structures within bone makes it challenging to translate SRH to bone imaging. More recently, we have shown that SRS imaging of chemical changes of mineral content of calcified breast tissue can also provide critical diagnostic information (*27*). In particular, lower carbonate levels of calcifications are strongly correlated with malignancy. We hypothesize that the chemical changes of bone composition are also reflective of underlying pathology, and SRS could potentially be used for intraoperative diagnosis of bone cancer.

This study aims to demonstrate the diagnostic utility of a multimodal approach to visualizing and analyzing the structure and chemical compositions of various bone specimens using SRS microscopy combined with two-photon fluorescence (TPF) and SHG. We used SRS to visualize unsectioned bone specimens containing both organic and mineralized components at high spatial resolution and chemical specificity. Because minimal or no processing of surgical specimens is needed, we were able to produce an unprecedented level of detail about bone in association with pathologies frequently encountered in intraoperative settings. A simple staining procedure and robust TPF process allowed us to highlight nuclear details to assist pathologists in extracting the relevant diagnostic information. In addition, we acquired SHG to evaluate collagen organization (*28*) as an additional metric of bone structure in different physiological and pathological conditions. Because SRS, TPF, and SHG share the same laser source and microscope, they can be acquired simultaneously, which greatly simplifies the imaging workflow and reduces imaging time. Our results show that chemical and morphological features obtained from the multimodal imaging method can distinguish specific categories of bone cancer with >90% accuracy and provide critical information needed for intraoperative consultation. Because our non-destructive imaging approach does not interfere with downstream histological and molecular analysis, it has the potential to fulfill a much-needed role in intraoperative diagnosis for orthopedic oncology.

## Results

### Imaging morphological and architectural features of bone specimen with SRS and TPF

One of the most useful diagnostic features of H&E is the morphology and architecture of cellular organization. For conventional H&E to work, bone must be demineralized and thinly sectioned. SRH does not require tissue processing. However, the same protein and lipid contrasts used in SRH for visualizing soft tissue do not generate sufficient contrasts for bone tissue (**Fig. S1**). An additional challenge is that un-sectioned bone tissue is highly scattering. It requires imaging in the epi-mode, instead of transmission mode which is employed in most SRH applications to date.

Here we demonstrate that using SRS imaging at ∼2930 cm^-1^ (a CH vibrational mode predominantly for protein) and TPF of acridine orange (*29*) we can visualize the morphology of bone and cellular structures within and nearby. Together SRS and TPF provide a simulacrum of the typical H&E, highlighting histologic features of bone tissue including cell type, stroma and matrix at high resolution. Both imaging modalities are implemented in the epi-mode (**Fig. S2**), which is amenable to surgical tissue of any size and shape with an imaging depth of ∼ 100 µm without additional processing.

Pathological bone tissues exhibit distinct morphological features. Chondrosarcoma is a malignant cartilage forming tumor that is a common bone malignancy and is presumed to arise from mesenchymal cells that differentiate along the chondrocytic lineage (*30*). Our case of chondrosarcoma (case 11 in **Table S1**) shows the characteristic gross appearance of cartilaginous neoplasm with glistening gray-white lobules (**Fig. 1A**, below the red dashed line) invading bone (**Fig. 1A**, above the red dashed line). Corresponding H&E (**Fig. 1B**) confirms the presence of polygonal to spindled neoplastic cells (green box in **Fig. 1B**) in a loose cartilaginous matrix invading adjacent cortical and medullary bone. The combination of SRS and TPF images are consistent with H&E, clearly showing neoplastic cells in a cartilaginous matrix invading bone (**Fig. 1C**). Our findings demonstrate that the integration of SRS/TPF imaging modalities allows for the visualization of cellular structures of neoplasm in bone specimens with high resolution, which correlates well with the histologic features seen in conventional H&E staining.

**Fig. 1.**
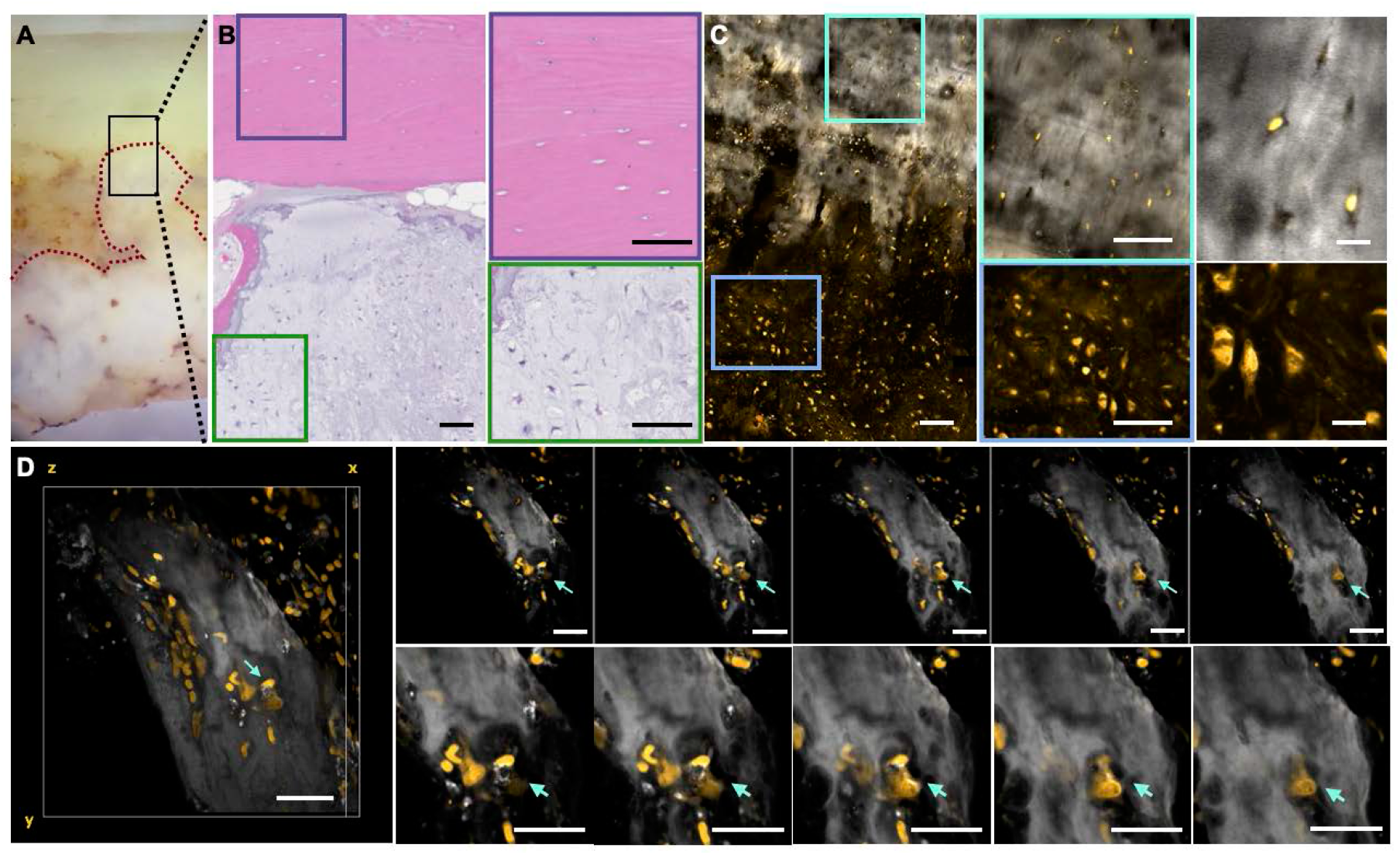
Imaging morphology of bone specimen. **(A), (B)** Gross image of bone adjacent to chondrosarcoma focus with corresponding H&E. Scale bars are 70 µm. **(C)** SRS/TPF imaging of the same geographical area (greys - SRS at ∼2930 cm^-1^ and gold – TPF from AO stained nuclei). Scale bars are 70 µm for larger view and two close ups (in blue and cyan). Scale bar is 20 µm for cellular features close up. **(D)** 3D stack of bone remodeled by osteoclasts as indicated by cyan arrows (the space between stacks is 2 µm).

In addition to being able to image a thick tissue specimen, SRS/TPF has an intrinsic capability to provide 3D information. This can overcome one of the drawbacks of conventional histology, where the information is mostly confined to the 2D plane. If a pathologist encounters a diagnostically challenging scenario where additional H&E sections can be beneficial, it is likely to take additional time for sampling, sectioning, and processing to produce additional tissue sections for evaluation if warranted during pathological examination. 3D imaging with SRS/TPF can be readily achieved in near real-time. The bone and its microenvironment are composed of various cell types. Our 3D reconstruction shows the presence of osteoblasts, the cell type responsible for bone formation, lining the bone trabeculae (**Fig. 1D**, indicated by a cyan arrow). We also observe cells within cavities, known as resorption pits. This morphology is consistent with osteoclasts. Similarly, pathologist can use SRS/TPF 2D and 3D capability to identify other cell types including osteocytes and chondrocytes (**Fig. S3**).

Overall, we show that SRS and TPF provide diagnostically useful morphological and architectural information that can assist a pathologist’s evaluation of bone specimens intraoperatively. The intrinsic optical sectioning of SRS and TPF enables 3D assessments of specimens when desired.

### SRS based chemical imaging of mineral content provides diagnostical information

In current pathology practice, one of the most underutilized diagnostic components of bone tissue is mineral content. Several publications suggest that the mineral component of bone is altered during cancer development (*19, 20, 31*). Here, we explore the diagnostic utility of chemical information provided by SRS in the fingerprint region as we interrogate unprocessed mineralized tissue involved in both non-neoplastic and neoplastic pathologic conditions.

**Fig. 2A-D** show an example of non-neoplastic bone sample. This case has a history of avascular necrosis (**Table S1**, case 5) and shows endochondral ossification, which occurs as a part of new bone formation. When imaged with SRS/TPF, multiple foci of newly formed bone were noted. A representative focus of osteoid is shown in **Fig. 2B-D** (indicated by star). SRS imaging at the hydroxyapatite peak ∼960 cm^-1^ (**Fig. 2C**) demonstrates gradual mineralization of the newly formed osteoid (indicated with pink star) as indicated by decreasing intensity of the image while moving from the area of more mature bone (indicated with pink arrow) to new bone. The corresponding carbonate content % map shows a low carbonate content in newly formed bone to be closer to 0 (**Fig. 2D**). The spectra (**Fig. 2E**) from those two distinct areas of endochondral ossification show the difference in carbonate/phosphate SRS intensity ratio. Our result is consistent with previous research showing that newly mineralized osteoid consists of predominantly hydroxyapatite (*32*).

**Fig. 2.**
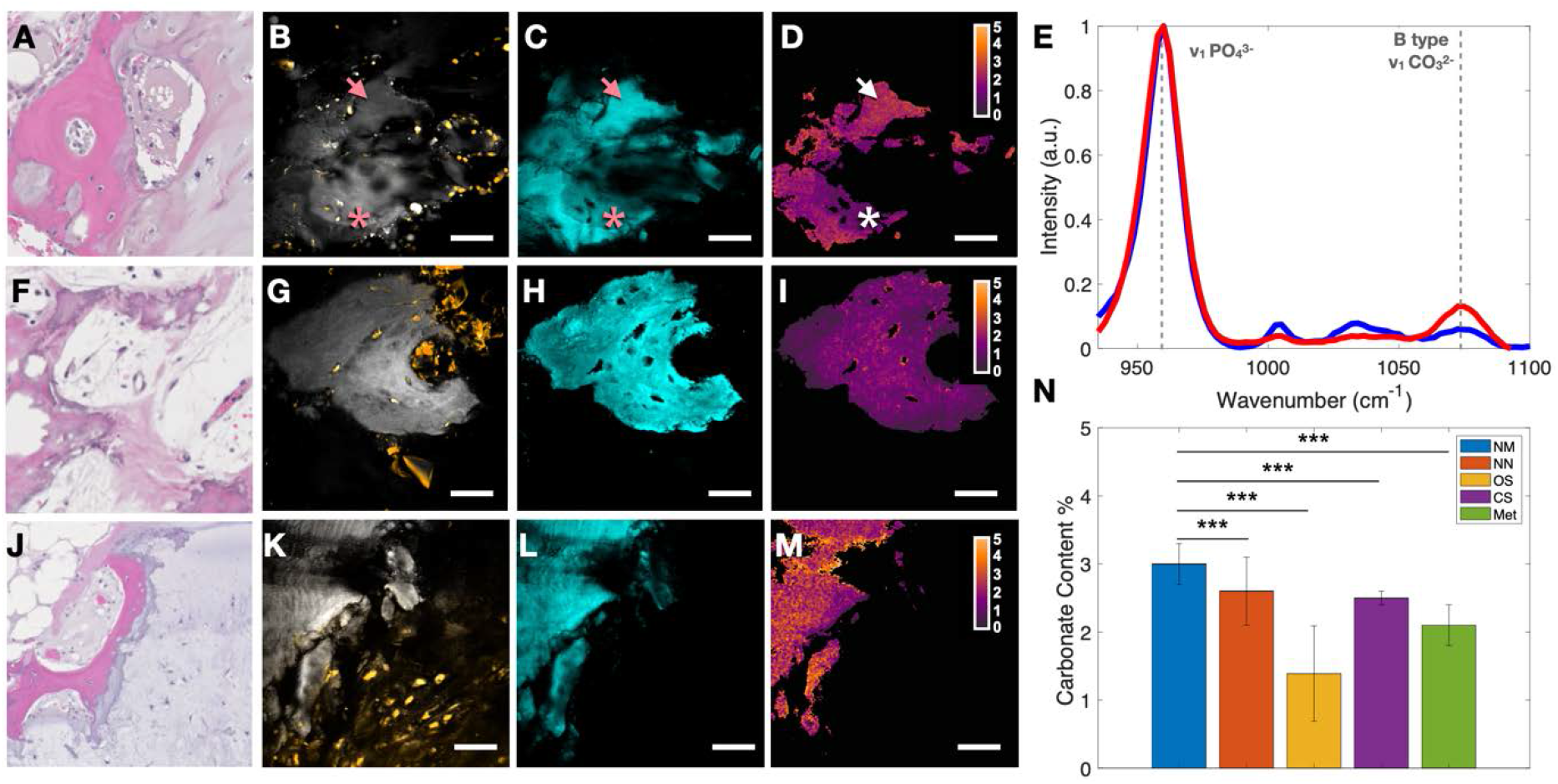
Chemical imaging of mineral bone component. **(A)** H&E of bone healing after avascular necrosis with endochondral ossification focus shown here. **(B)**-**(D)** Representative area showing new bone formation with contrasts from morphology (B, greys - SRS at ∼2930 cm^-1^, gold – TPF from AO stained nuclei collagen), mineralization (C, cyan – SRS at ∼960 cm^-1^), and mineral content (D, carbonate level), respectively. Pink/white arrow corresponds to the mature bone. Pink/white star corresponds to osteoid forming new bone. **(E)** Spectra of two different areas identified in **(D)**. White arrow corresponds to the spectrum in red. White star corresponds to spectrum in blue. **(F)** H&E of osteosarcoma case with the residual neoplastic bone matrix. **(G)**-**(I)** Representative area showing neoplastic bone matrix (greys - SRS at ∼2930 cm^-1^, gold – TPF from AO stained nuclei collagen, cyan – SRS at ∼960 cm^-1^, and SRS derived carbonate content map). **(J)** H&E of bone involved by chondrosarcoma. **(K)**-**(M)** Representative area showing bone adjacent to neoplastic cells (greys - SRS at ∼2930 cm^-1^, gold – TPF from AO stained nuclei collagen, cyan – SRS at ∼960 cm^-1^, and SRS derived carbonate content map). **(N)** Bar chart of carbonate content % across groups imaged in this study (NM-normal bone, NN-non-neoplastic pathological process including bone remodeling, OS – osteosarcoma, CS – chondrosarcoma, Met – metastatic cancer). Two-sided t-test is performed and *** - p-value<0.001. Error bar is represented by standard deviation.

In contrast, the analysis of carbonate content in a case of osteosarcoma (**Table S1**, case 9; representative area shown in **Fig. 2F-I**) shows that the entire neoplastic bone matrix has low carbonate content. This is likely attributed to the higher acidity of the cancer microenvironment precluding the inclusion of carbonate ions (*33*). Our result suggests that neoplastic bone growth is distinct from normal bone growth, where mature bone has substantially higher carbonate content.

Interestingly, the carbonate level and heterogeneity also reflect the type of neoplastic process. For comparison, we show that the bone adjacent to the neoplastic cells in a case of chondrosarcoma (**Table S1**, case 11) mostly retains high carbonate mineral content (representative area in **Fig. 2J-M**). Only regions close to the neoplastic cells appears to be impacted by the cancer microenvironment displaying lower carbonate content. Chondrosarcoma does not produce osteoid and thus does not affect mature bone that is already present. Thus, the observed spatial gradient of carbonate content is likely a direct result of the underlying neoplastic process.

To evaluate the variations in carbonate content for different pathological conditions, we compared the average carbonate content across all groups of non-neoplastic and neoplastic cases included in this study (**Fig. 2N**). The non-neoplastic pathologic conditions are avascular necrosis, abnormal bone hypertrophy, and fracture. The neoplastic samples analyzed are primary bone neoplasms (osteosarcoma and chondrosarcoma) and metastatic cancer to bone. All pathologies show statistically significant difference from the normal. The osteosarcoma cases appear to have the lowest carbonate content relative to normal and other groups. Our results confirm our hypothesis that chemical changes of bone composition reflect underlying pathology and can be used to differentiate neoplastic cases.

### Organic matrix changes are indicative of bone pathology

The organic matrix of bone is primarily composed of type I collagen and is typically formed by osteoblasts. Newly formed bone matrix, osteoid, lacks mineralization. When osteoblasts are surrounded by osteoid, they differentiate into osteocytes. The lacunae, spaces where osteocytes remain for the duration of their lifetime, vary in shape and size. Lacunae are typically round to almond-shaped and then often become more oblong as they mature (*34*). Because bone formation, remodeling, and repair are dynamic processes responsive to the microenvironment, we hypothesize that morphological changes of lacunae and collagen are indicative of underlying pathological processes and can be used as diagnostic parameters.

To test our hypothesis, we performed morphometric analysis of osteocyte lacunae for different pathological conditions (**Fig. 3A**). Several parameters of interests for lacunae and collagen can be determined from SRS/SHG imaging, including lacuna area, aspect ratio (AR, defined as the ratio between the major axis and the minor axis of a lacuna), collagen anisotropy (*35*), and the angle between lacuna and collagen. The lacunae in the SRS image appeared dark and were hand segmented for this purpose (**Fig. 3B)**. Although we primarily used manual segmentation here, the automation of such a process is possible using machine learning platforms such as Ilastik (*36*). Using particle analysis available in ImageJ, we obtained aspect ratios of segmented lacunae as well as their areas (A). Additionally, the angle of the major axis along a lacuna relative to the horizontal (β_L_) is determined. We used SHG from collagen to measure matrix organization (**Fig. 3C**). Using FibrilTool (*35*), we determined the average angle of collagen relative to horizontal (β_C_) and anisotropy value (Φ) of the collagen fibers. For anisotropy assessment with Φ of 1 representing complete co-alignment and Φ of 0 representing complete lack of co-alignment. To generate a map of β_C_ and Φ, an in-house algorithm was used to apply FibrilTool to 64 × 64 pixels^2^ regions while scanning across the image (**Fig. 3D-E**, see methods section for additional detail). The area of lamellar growth highlighted with SHG corresponds to Φ approaching 0.5. In comparison, a basketweave pattern of collagen has lower anisotropy. Such difference highlights the potential to identify pathological conditions where collagen growth is impaired or significantly different.

**Fig. 3.**
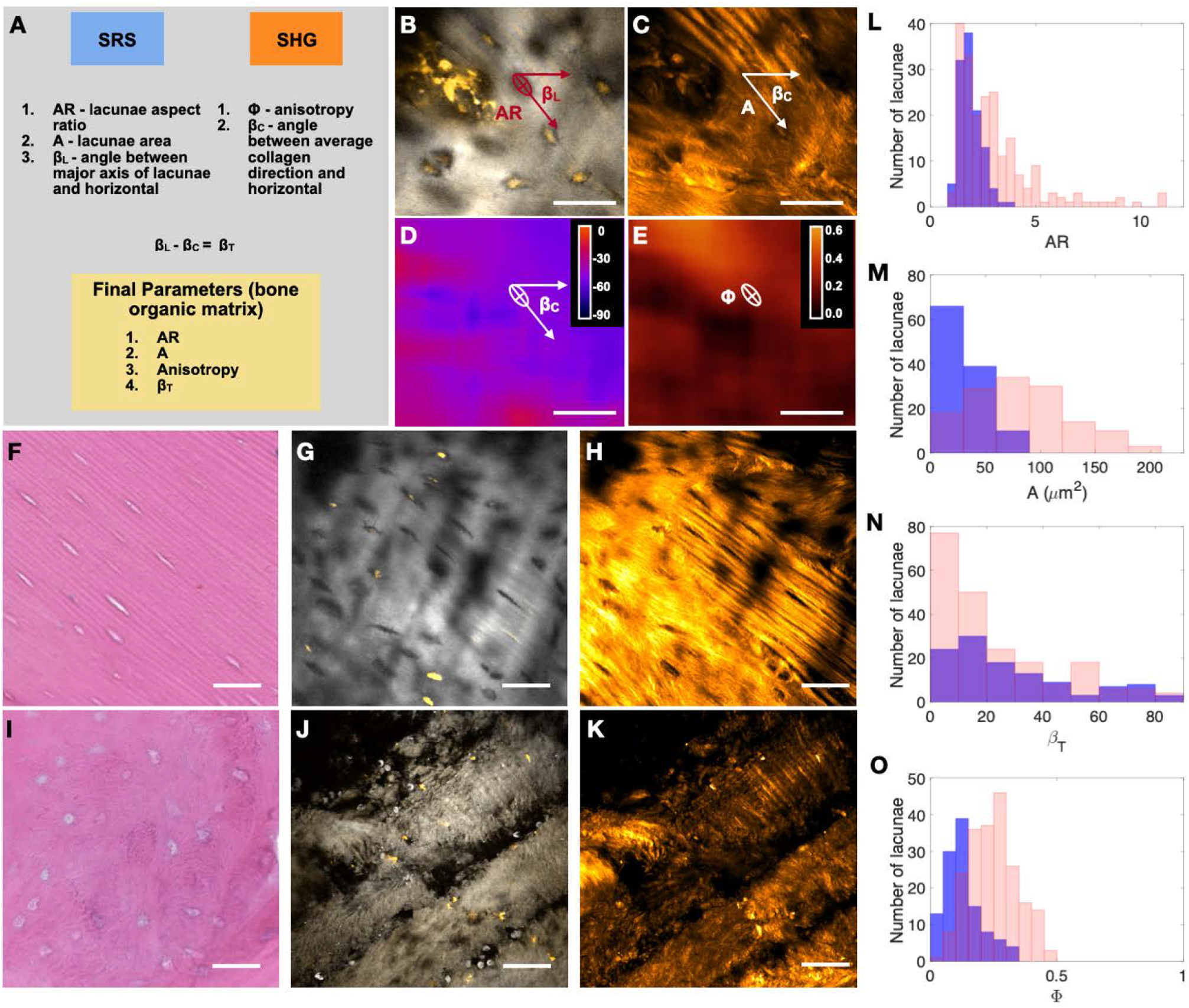
Morphometric analysis of organic bone matrix with examples highlighting the differences in the organic bone matrix as determined by morphometric analysis and supported by histology. **(A)** Depiction of parameters used for morphometric analysis. **(B)** Lacunae aspect ratio AR and angle for major axis of lacunae with horizontal β_L_ are determined from SRS (2930 cm^-1^) data. **(C)** Angle of collagen with horizontal β_C_ and collagen anisotropy A are determined from SHG using FibrilTool. Parameter β_C_ is defined as an angle collagen makes with horizontal. **(D), (E)** Resulting angle of collagen with horizontal and anisotropy maps used by ImageJ ROI manager to determine β_C_ and Φ associated with a given lacuna (highlighted with ellipse). **(F)**-**(H)** H&E, SRS/TPF, and SHG images of normal bone. **(I)**-**(K)** H&E, SRS/TPF and SHG images of hypertrophic bone. **(L)**-**(O)** Histograms for aspect ratio of lacunae AR, lacunae area A, angle of nearby collagen, and the major axis of lacunae β_T_, and anisotropy as determined by FibrilTool for normal bone (red) and hypertrophic bone (blue).

When comparing normal mandible and hypertrophic mandible from abnormal development, hypertrophic mandible shows a much more disorganized collagen morphology. In addition, its lacunae are smaller and rounder (**Fig. 3F-K, Fig. S4** with confirmatory H&E). From the histogram, we can observe that normal bone has a broader distribution of AR values and lacunae sizes compared to hypertrophic bone (**Fig. 3L-M**). Normal bone generally has larger osteocyte lacunae (**Fig. 3M**) with better coaligned lacunae and nearby collagen (quantified by coalignment angle β_T_ = |βc-β_L_|, as shown in **Fig. 3N**). In contrast, in abnormal bone, the coalignment angle has a broader distribution. Collagen also has a higher degree of disorganization (lower anisotropy Φ) in abnormal bone compared to normal bone (**Fig. 3O**).

Applying the same analysis across all samples, we summarized the differences in morphological parameters among different pathological conditions (**Fig. 4**). The aspect ratio of lacunae across all pathological categories shows the most differences between normal and non-neoplastic pathology as well as metastatic cancer (**Fig. 4A**). These findings are consistent with visual observations where lacunae in osteoid or newly formed bone appear to have lower aspect ratio relative to mature bone. Moreover, the area of lacunae is variable across all groups with the most significant differences occurring between normal cases and cases that include healing bone and neoplastic bone matrix (**Fig. 4B**). When evaluating collagen organization, normal bone and bone with metastatic cancer show the largest difference in the coalignment angle (β_T_) (**Fig. 4C**) and anisotropy (Φ) (**Fig. 4D**). It is plausible that these parameters are capturing higher bone turnover and subsequent disorganization of the matrix in metastatic cases (*37*).

**Fig. 4.**
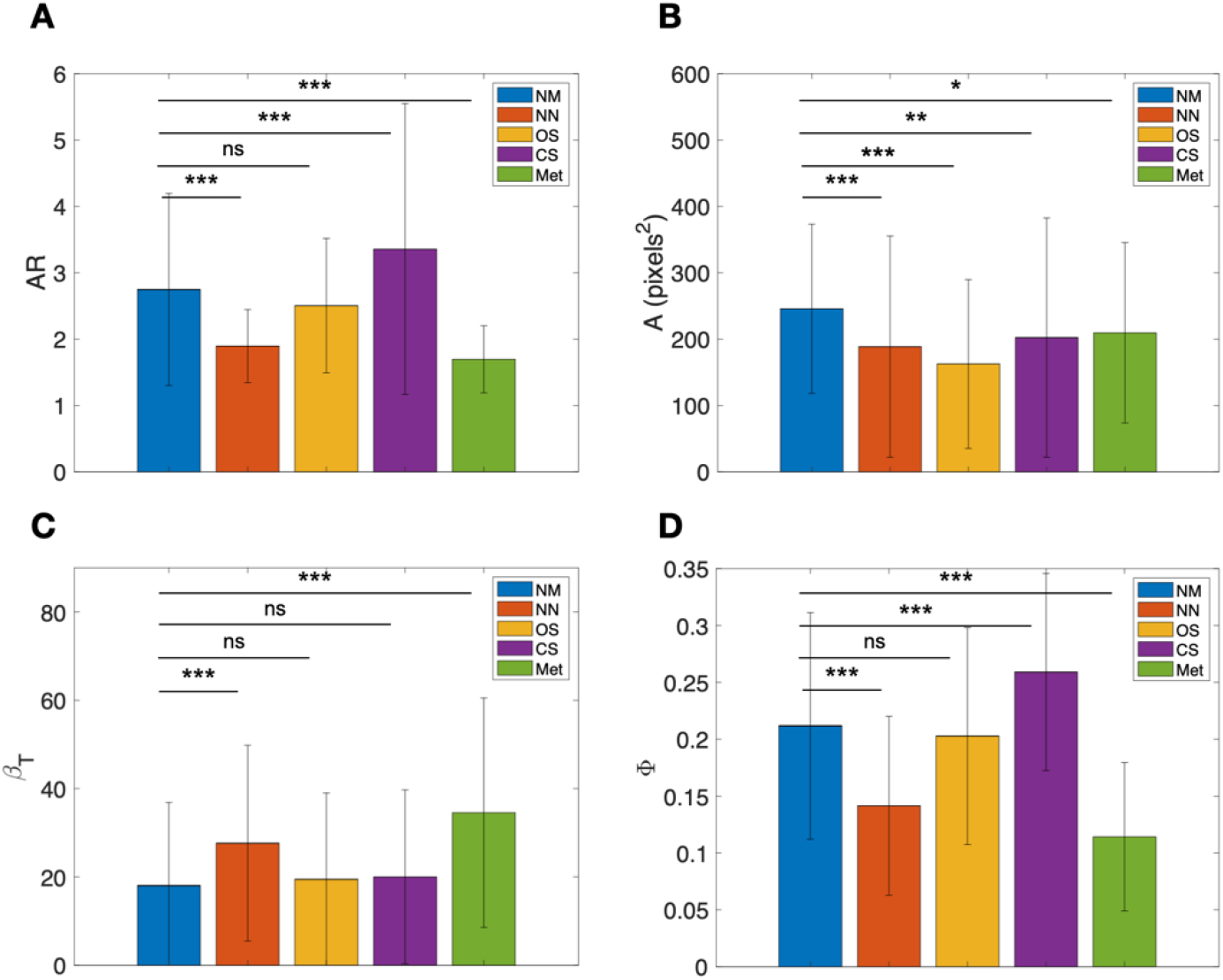
Summary of morphometric analysis for bone specimens. **(A)** Bar chart for aspect ratio of lacune AR averages. **(B)** Bar chart for lacunae area A. **(C)** Bar chart for angle of nearby collagen and major axis of lacunae β_T_. **(D)** Bar chart for anisotropy Φ. Error bar is reflective of standard deviation. Two-sided t-test is performed and * denotes p-value < 0.05, ** – p-value <0.01, and *** - p-value<0.001. NM-normal bone, NN-non-neoplastic pathological process including bone remodeling, OS – osteosarcoma, CS – chondrosarcoma, Met – metastatic cancer. N_NM_ = 274 lacunae, N_NN_ = 178 lacunae, N_OS_ = 119 lacunae, N_CS_ = 114 lacunae, N_Met_ = 106 lacunae.

### Combined morphometric and chemical analysis enables highly accurate classification of bone pathologies

As shown in the previous sections, both mineral components and morphometric analysis of lacunae and collagen matrix provide diagnostically useful information. In an intraoperative setting, it is important to determine if the lesion is benign or malignant and in the case of malignant, further subdivide into primary and metastatic disease. We demonstrate that by combining chemical and morphological analysis enabled by multimodal imaging, we can achieve highly accurate classification of specific bone cancer types. To build a robust classification algorithm with multiple parameters, we employed a supervised machine learning model. The advantage of a machine learning algorithm is that once a model is trained, predictions can be generated in almost real time which can help to guide intraoperative pathological diagnosis and subsequent patient treatment. We performed Random Forest classification using TreeBagger in Matlab. It is a bootstrap-aggregated (bagged) decision tree where multiple decision trees are created using randomized subsets of the data and features from the training set for each tree. The ensemble of trees can then be applied to new data where each tree “votes” on the correct prediction where the final category is determined by the greatest number of votes. By selecting a random subset of predictors for the generation of each tree, the algorithm is able to reduce the effects of overfitting and improves generalization (*38*).

We combined the data from the morphometric analysis (including aspect ratio, area of lacunae, nearby collagen anisotropy, and the coalignment angle between collagen and lacunae central axis) with carbonate content for each lacuna. Using these parameters for 791 lacunas, we constructed a classification model using a training data set consisting of randomly selected observations from the full dataset (70% of the original data) and retained a separate validation data set consisting of the remaining data (30% of the data).

By determining the minimum number of trees needed to reach an adequately reduced Out-of-Bag (OOB) classification error (**Fig. 5A**), we minimize the possibility of overfitting the data. The final number of trees that was chosen for our classification model was 15. There was no significant difference in error when considering more than 15 trees.

**Fig. 5.**
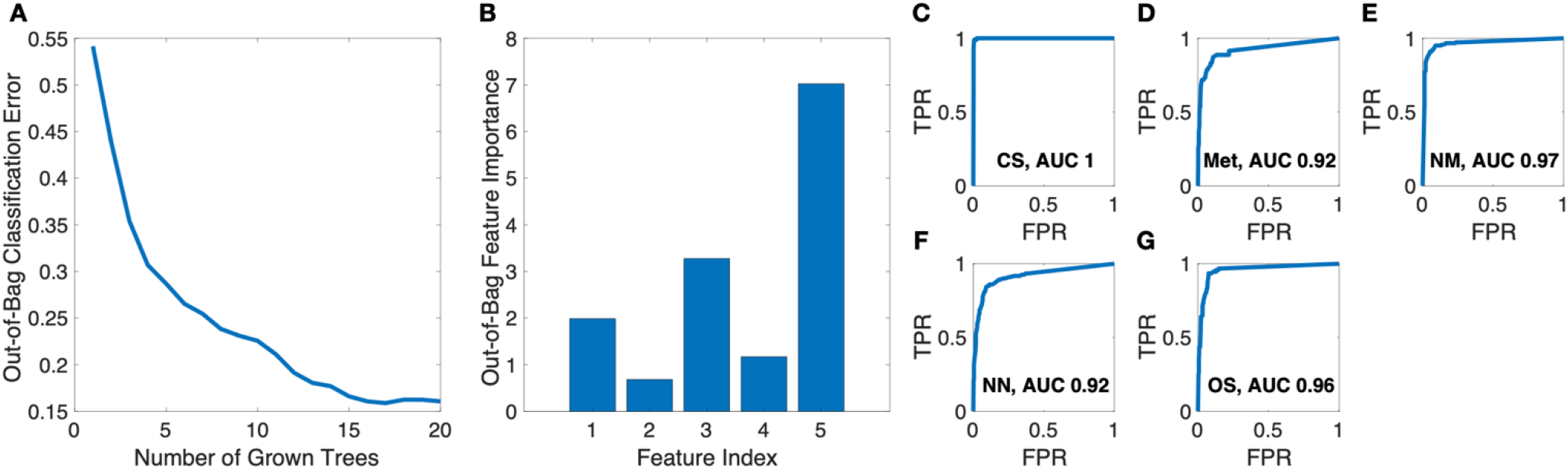
Bootstrap-aggregated (bagged) decision tree-based classification model using parameters from organic matrix morphometric analysis in conjunction with carbonate content in bone mineral matrix. **(A)** Out-of-Bag (OOB) classification error vs number of grown trees. **(B)** Out-of-Bag features importance *versus* features index (1 - AR, 2 - Φ, 3 - β_T_, 4-A, 5 - carbonate content of mineral portion of bone). **(C)**-**(G)** ROC curves for diagnostic groups used in this study (NM-normal bone, NN-non-neoplastic pathological process including bone remodeling, OS – osteosarcoma, CS – chondrosarcoma, Met – metastatic cancer). TPR – true positive rate. FPR – false positive rate.

Based on the results of OOB feature importance assessment (**Fig. 5B**), AR, anisotropy, and the angle between collagen and lacunae central axis contribute similar weight to final decisions. The area is less important, probably due to that fact that SRS/TPF imaging has optical sectioning effect. Depending on the location of the sectioning, the area can be quite variable and thus not reliable for differentiation. The carbonate content percentage as determined by SRS imaging is most indicative of the underlying pathologic bone condition. The results of the model built without carbonate data is included in supplemental (**Fig. S5**). This is consistent with our observation that carbonate level is significantly different among various pathological conditions, even without spatially resolved information.

The receiver operating characteristic (ROC) curve is used to evaluate the model effectiveness for identifying a particular diagnosis among different diagnostic entities including normal bone, non-neoplastic pathologic bone conditions, primary malignant bone tumors and metastatic cancer to bone (results in **Fig. 5C-G**). The area under the curve (AUC) of 0.8 to 0.9 is considered excellent and above 0.9 is considered outstanding in medical practice (*39*). The areas under the curve (AUCs) were calculated to determine how well a given diagnostic group can be separated from others. Overall accuracy of predictions was calculated by comparing known diagnostic category with model-predicted category and resulted to be ∼90 ± 2%.

In summary, the results of the classification model show that lacunae associated parameters obtained by morphometric analysis in conjunction with carbonate content for the mineral portion of bone can provide a good basis for assessing bone specimens in an intraoperative setting.

## Discussion

Interrogation of bone specimens is severely limited by conventional histological methods due to their reliance on sectioning of tissue into 4-6 µm sections. Without the ability to visualize pathological processes of bone tissue, intraoperative consultations are hindered and can rely only on soft tissue, if present. These limitations often lead to difficulties in surgical decisions and the need for multiple surgeries in many cases. In this paper, we conduct the first-in-class study to demonstrate how multimodal non-linear optical imaging have a remarkable potential to provide a much-needed tool to help facilitate diagnostic classification and clinical decisions.

By using multimodal imaging, we can visualize bone structure and chemistry in a non-destructive manner. Being able to provide such information with relative ease and without compromising the availability of tissue is important in the context of an intraoperative setting as it preserves the tissue for subsequent molecular studies if they are deemed necessary. TPF from acridine orange and SRS of proteins allow us to generate images that could be used similarly to those from H&E. Using distinct Raman features of mineral components, particularly phosphate and carbonate, we can probe mineral components of bone and study how changes in mineral content correlate with bone pathology, including neoplastic processes. By collecting SHG from collagen fibers, we augment collected data with important information about surrounding collagen fiber organization.

Our main findings show that carbonate content varies across different pathological categories, particularly in osteosarcoma. Being able to diagnose osteosarcoma and separate this entity from fractures or other processes that do not warrant complete excision offers much-needed actionable information to the surgeon. Additionally, morphometric analysis of bone organic matrix provides additional parameters (lacunae aspect ratio, angle of collagen relative to lacunae major axes, and collagen anisotropy) that prove to be useful in identifying cases where the organic matrix is abnormal, specifically in cases of abnormal bone healing/growth and metastatic cancer. In the cases where excision is warranted, non-destructive imaging in combination with analysis can potentially provide diagnostic information necessary for margin assessment. By combining morphological and chemical features with a machine learning approach (Random Forest classification), we demonstrate that an overall accuracy of >90% can be achieved in classifying specific classes of pathology including non-neoplastic condition, primary malignant bone tumors (osteosarcoma and chondrosarcoma), and metastatic cancer to bone. Our results suggest that multimodal nonlinear optical imaging has the potential to address the major gap in intraoperative pathology involving bony specimens.

One limitation of the current study is the sample size. A larger study involving many more patients will be necessary to fully validate the method for bone cancer classification and develop a robust algorithm for clinical applications. In an intraoperative setting, it is necessary to have miniatured laser and rugged microscope without stringent environmental control. Such efforts are already underway for brain cancer surgery (*22*). The other limitation is data acquisition speed. With multimodal imaging, multiple contrasts are generated simultaneously. However, the hyperspectral SRS imaging takes extra time due to the acquisition at multiple Raman transition regions. Excessive time on the order of many hours is still needed for imaging large tissue samples (cm in size) at the moment. The multimodal and hyperspectral imaging approach generates a large swath of spatial-spectral features and we only use part of the data available in the analysis. Unbiased and robust diagnosis will benefit from advances in machine learning approaches that deal with large datasets, particularly deep learning, when large-scale testing is available (*24, 25, 40*). Machine learning will also help extract parameters that are most important for diagnosis and minimize the amount of data required, thus shortening the data acquisition time in an intraoperative setting. Ultimately, combining advances in multimodal non-linear imaging modalities and machine learning, we can potentially enable surgeons and pathologists to quickly determine the correct diagnosis and necessary treatment, as well as reducing unnecessary surgery and delay in getting the final diagnosis, especially in orthopedic oncology.

## Materials and Methods

### Multimodal imaging with SRS, TPF, and SHG

The details for the broadband SRS set up were outlined in previous publications (*27, 41*). Briefly, we used broadband femtosecond dual-beam laser system (Insight DS+ from Spectra-Physics) where a tunable beam (pump) was centered at 798 nm for CH Raman region and 944 nm for fingerprint Raman region and a fixed beam (Stokes) was centered at 1040 nm. A home-built laser scanning microscope equipped with a 25× Olympus water immersion objective (NA=1.05) was used to image the sample (**Fig. S2**). The SRS signal is detected in epi mode with a large area silicon photodiode, while epi-fluorescence from TPF and SHG are detected simultaneously by a photo-multiplier tube (Hamamatsu H10770PA-40). At the focus, the pump and Stokes beams had an average power of 50 mW each. For SRS imaging, a stack of frames with single field of view (FOV) of 285 μm × 285 μm (512 pixels × 512 pixels) was acquired, for which the pixel dwell time is 8 μs and time to acquire each frame is 2 s. The frames are acquired every 2 cm^-1^ for total of 90 frames on average. For SHG and TPF imaging a bandpass filter (520 nm +/- 20 nm) was used.

Bone specimen is placed into an aluminum sample holder covered with a glass coverslip (**Fig. S6A**). The tissue was photographed to map imaged areas. For each sample, an area of ∼2 × 2 mm at the bone-soft tissue border was imaged with SRS at ∼2930 cm^-1^ corresponding to the protein vibrational peak, TPF from acridine orange stained nuclei and SHG from collagen (**Fig. S6B-F**). Detailed SRS data was collected in both high-wavenumber and fingerprint Raman regions as hyperspectral stacks in selected regions of interest (ROI). The high-wavenumber SRS images on C-H stretching were used to determine lipid and protein components in studied tissues. The SRS fingerprint images were used to determine mineralization species present in our samples.

### Bone tissue specimens

Bone tissue samples was obtained from 19 patient cases with IRB approval from the University of Washington. The cases were de-identified and curated by Northwest Biotrust Biorepository. The case selection was based on the pathology report to include normal (rib, mandible, femoral head), non-neoplastic pathology (avascular necrosis, fracture, hypertrophic bone), primary neoplastic (osteosarcoma and chondrosarcoma), and secondary neoplastic (metastatic cancer to bone). **Table S1** includes additional details of the samples. The bone tissue was cut into 2-4 mm sections with a band saw and fixed in 10% neutrally buffered formalin. Two cases with sarcoma and metastatic to bone lung cancer were excluded due to inadequate bone available in samples.

### Tissue staining

Before imaging, bone tissue was stained with acridine orange (5 μg/mL as dissolved in Dulbecco’s phosphate buffered saline). The bone tissue was stained for ∼10 min and thoroughly washed in phosphate buffered saline before imaging.

### Agar gel preparation

For imaging, bone tissue was imbedded into agar gel to immobilize the specimen. Agar gel (15 mg agarose/mL) was prepared by mixing 125 mmol/L NaCl, 10 mmol/L glucose, 10 mmol/L HEPES, 3.1 mmol/L CaCl_2_, and 1.3 mmol/L MgCl_2_ with agar (Sigma Aldrich) (*42*). The mixture was heated to 60 °C to liquify the gel for the embedding procedure.

### Sample mapping

Gross photographs were obtained of each specimen. Before imaging each sample, the sample stage position was recorded to determine the dimensions of sample and positions of main lesional area and subareas for imaging. Additionally, the depth of images was measured relative to the cover slip surface. The recorded locations of imaged tissue within the sample was correlated with histology after the experiment to confirm diagnosis and establish microenvironment for imaged bone.

### Calibration of SRS imaging to determine carbonate content of bone mineral conten

Chemical imaging and carbonate content determination was previously described (*27*). Briefly, calcium hydroxyapatite (HAP) and 10% carbonated hydroxyapatite (CHAP) were obtained from Sigma-Aldrich and Clarkson Chromatography Products respectively. Grounded powders were mixed and prepared for calibration (0%, 2.5%, 5.0%, 7.5%, 10% carbonate content). To determine carbonate content in bone samples (**Fig. S6H**), we used ratiometric SRS images at (∼1070 cm^-1^) and (∼960 cm^-1^) (*27*).

### Morphometric analysis of organic bone matrix

Using a combination of manual selection and interactive machine learning and segmentation toolkit, Ilastik (*36*), the osteocyte lacunae were segmented and stored as individual regions of interest (ROIs). Using Particle Analyze in ImageJ, the lacunae aspect ratio, area, and angle of major axis relative to horizontal were determined. Collagen organization was interrogated through SHG images. Using ImageJ FibrilTool (*35*) and in-house algorithms written in Matlab and ImageJ, we determined the angle of lacunae relative to collagen as well as local anisotropy. To generate a map, a sliding box (64 × 64 pixels) with sampling every 8 pixels was used. Using the ROI manager in ImageJ, angle and anisotropy data were determined for each individual lacuna.

### Statistical analysis

Using the parameters (aspect ratio, lacunae area, anisotropy, and angle) from the morphometric analysis and the carbonate content % obtained from SRS fingerprint data, a Random Forest classification model was generated. Specifically, we employed the TreeBagger algorithm in Matlab. We used 70% of original data for training and the remaining 30% used for testing the final accuracy.

## General

The authors of the paper especially appreciate the work done in support of our work by Marlie Reinmuth, Piper Driskell, and Sarah Bowell from Northwest Biotrust, Seattle, WA.

## Author contributions

D.F. and E.C. conceived and designed the project. K.S. performed most of the sample preparation, data acquisition and data analysis. E.C. supervised the tissue procurement and pathology analysis. D.F. supervised image acquisition and image analysis. S.M., A.W., and C.C. assisted with sample preparation, data acquisition, and data analysis. K.S., E.C., and D.F. wrote up the manuscript.

## Data and materials availability

The tissue samples were procured and de-identified by Northwest Biotrust, Seattle, WA. Data are available upon reasonable request.

