## Supplemental figures and tables for "Non-destructive Chemical Imaging of Bone Tissue for Intraoperative and Diagnostic Applications"

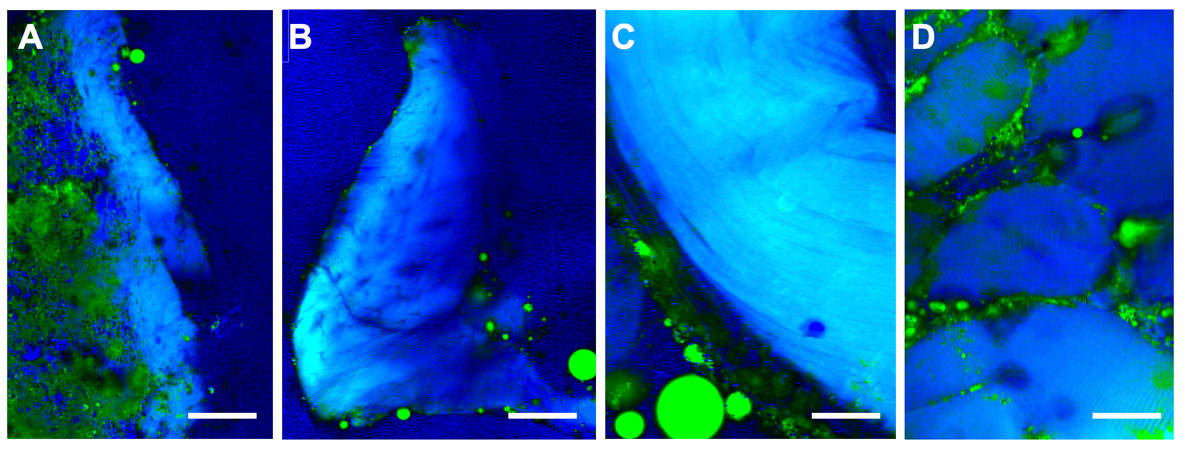

**Fig. S1. SRH of protein rich tissues with protein image in blue and lipid image in green. (A), (B)** SRH images of bone fragments with easily identifiable bone marrow elements in **(A)** and lacunae in **(B)**. Scale bar is 50 µm. **(C)**, **(D)** SRH images of tendon and muscle fragments. Scale bar is 40 µm for **(C)**. Scale bar is 100 µm **(D)**.

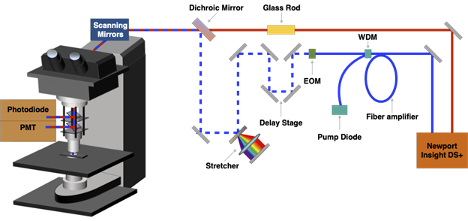

**Fig. S2. Schematic representation of SRS and SHG microscopy experimental setup.** EOM: electro-optic modulator. PMT: photomultiplier tube. WDM: wavelength division multiplexer.

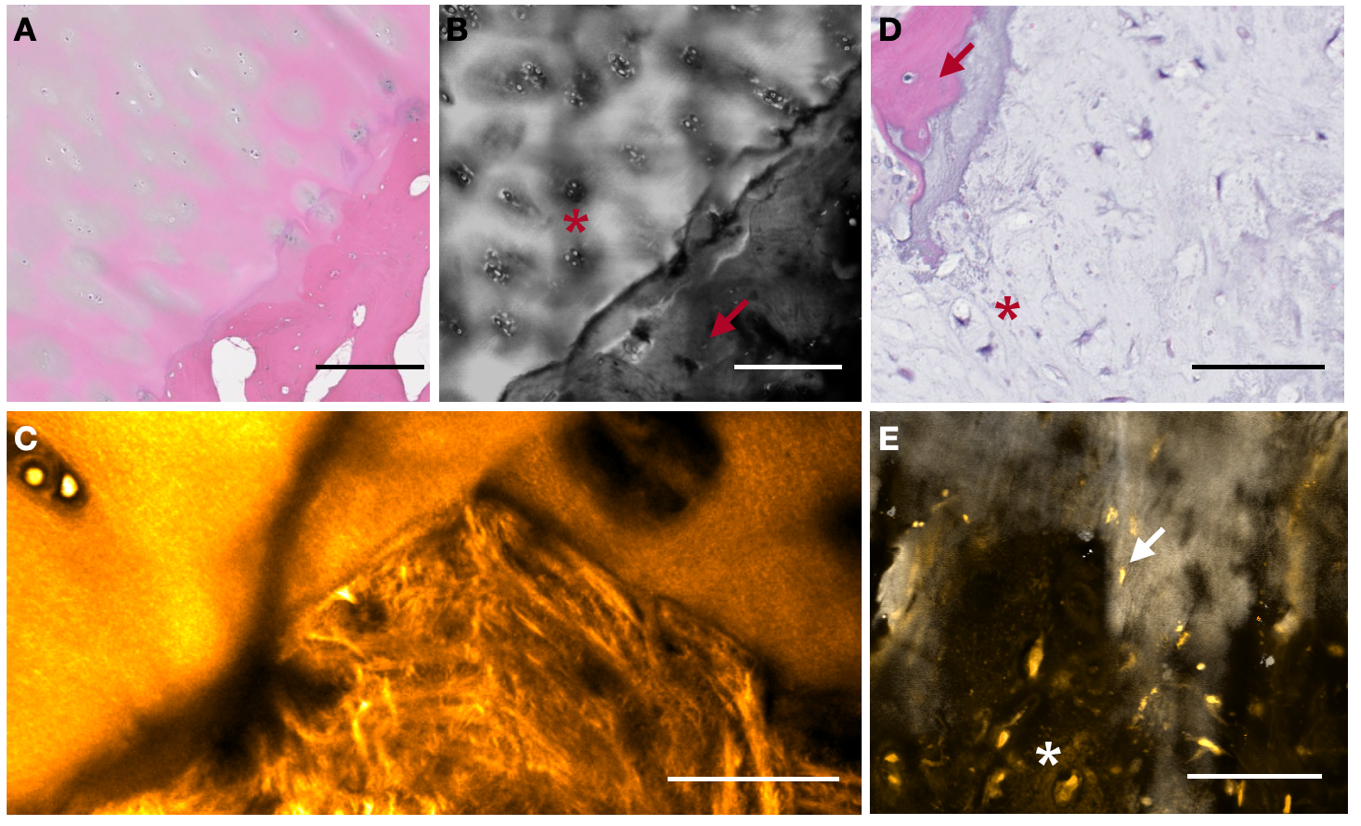

**Fig. S3. Chondrocytes and osteocytes with distinctly different matrix.** **(A)** H&E of femoral head highlighting cartilage overlaying bone. Scale bar is 250 µm. **(B)** SRS protein showing a clear border between cartilage and bone. Chondrocytes and osteocytes are identified with red star and red arrow, respectively. Scale bar is 250 µm. **(C)** SHG close up showing two distinct collagen organization patterns for cartilage and bone matrix. Scale bar is 50 µm. **(D)**, **(E)** Chondrosarcoma and nearby bone with identified chondrocytes and osteocytes. Scale bar is 70 µm.

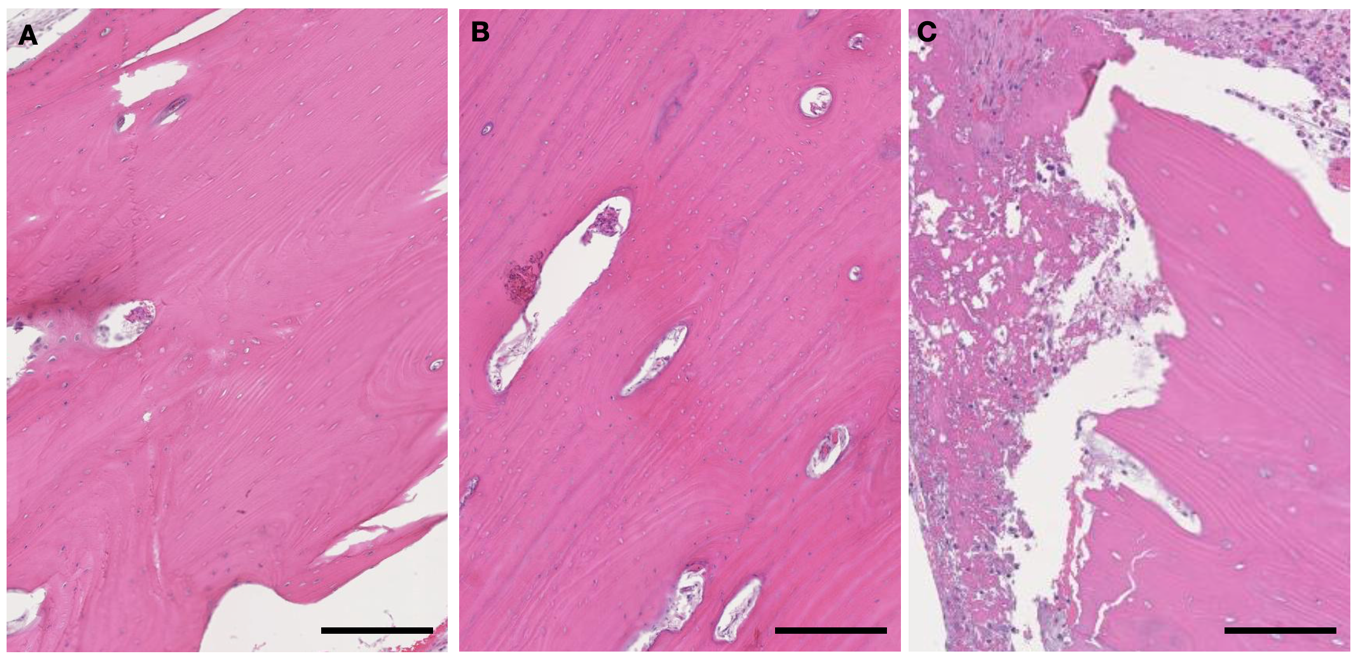

**Fig. S4. H&E for selected bone samples.** **(A)** Normal bone. **(B)** Hypertrophic bone. **(C)** Bone with nearby metastatic cancer. Scale bar is 250 µm.

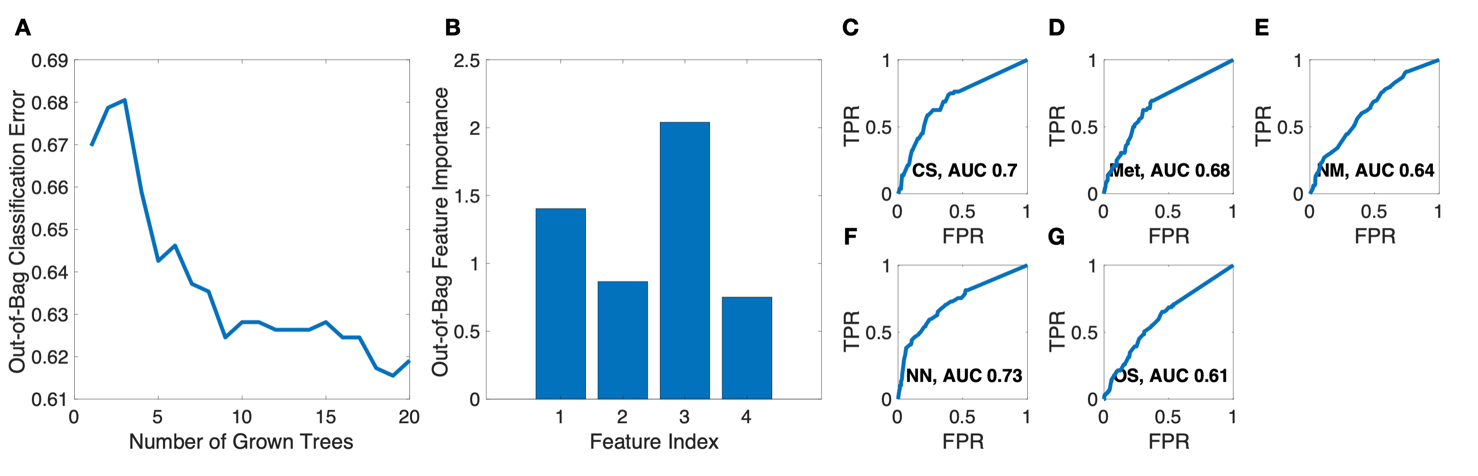

**Fig. S5. Bootstrap-aggregated (bagged) decision tree-based classification model using parameters from organic matrix morphometric analysis only.** **(A)** Out-of-Bag (OOB) classification error vs number of grown trees. **(B)** Out-of-Bag features importance *versus* features index (1 - AR, 2 - Φ, 3 - β_T_, 4-A). **(C)**-**(G)** ROC curves for diagnostic groups used in this study (NM-normal bone, NN- non-neoplastic pathological process including bone remodeling, OS – osteosarcoma, CS – chondrosarcoma, Met – metastatic cancer).

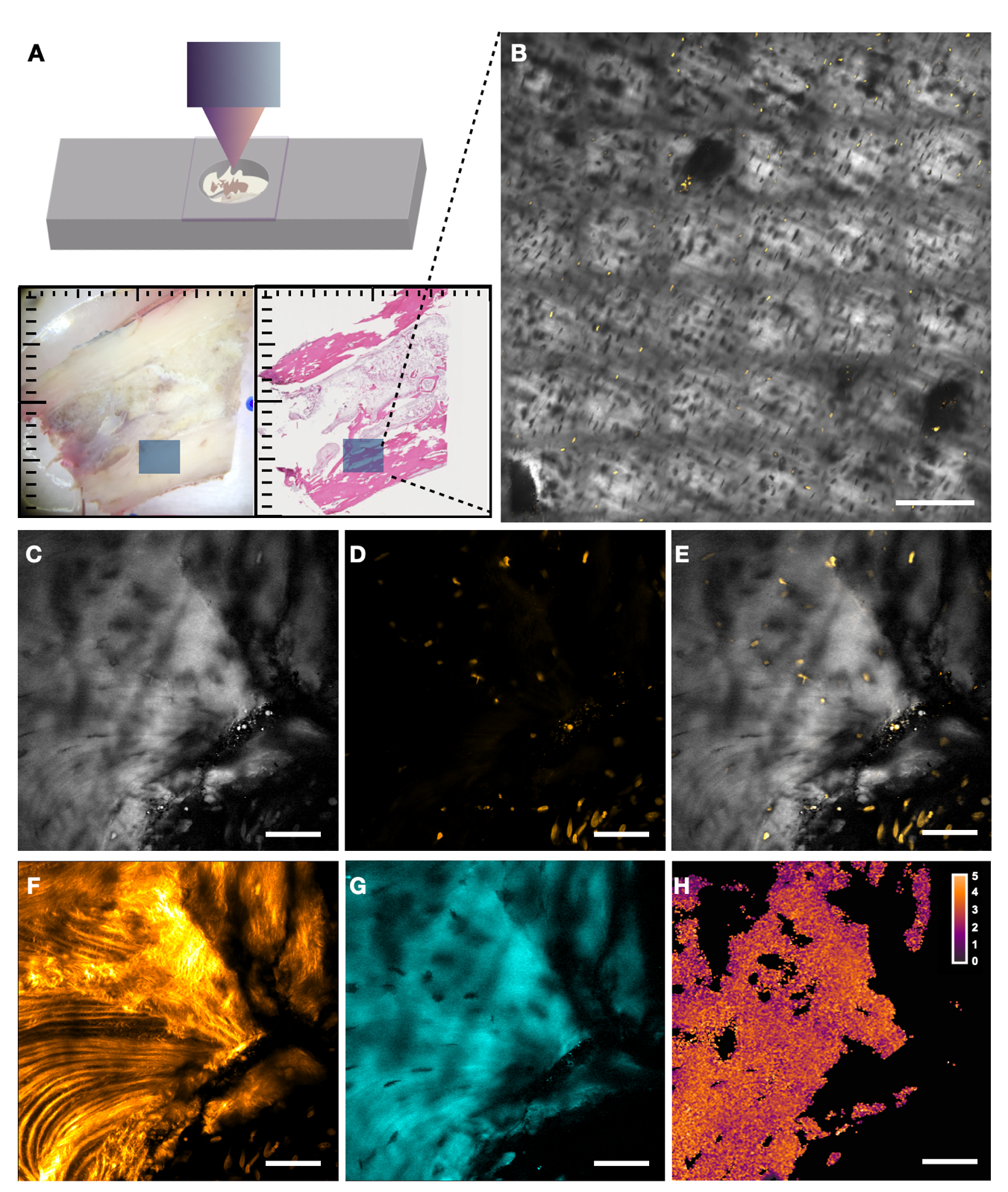

**Fig. S6. Workflow for bone tissue imaging and image processing.** **(A)** Graphical representation of a bone sample embedded in agar gel and sample holder. Gross image (left) in addition to post-imaging histology (right) are included. **(B)** An area to visualize bone microenvironment provides overview of tissue from where subareas are selected for more detailed study using hyperspectral microscopy. Scale bar is 250 µm. **(C)**-**(E)** Images obtained from SRS at ~2930 cm^-1^ (grey), TPF from AO stained nuclei (gold), and combination of both respectively. **(F)** SHG image obtained to highlight collagen organization. **(G)** SRS image at 960 cm^-1^ to highlight mineral content (cyan). **(H)** Carbonate content % map generated using fingerprint information from SRS (see Methods section). **(C)**-**(H)** scale bar is 50 µm.

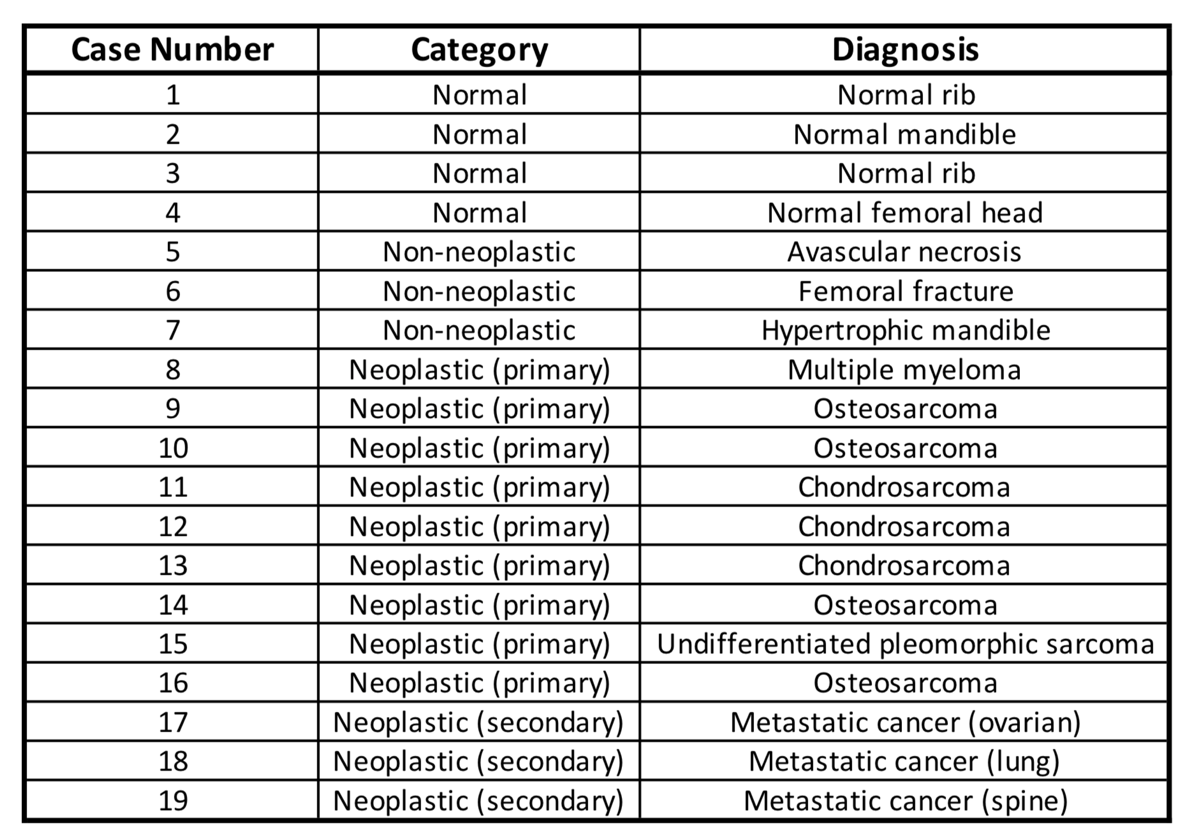

**Table S1. Description of case categories and diagnoses collected in the course of this study.**
